# Increasing Applicability of Automated mD-LC-MS Peptide Mapping for Biopharmaceuticals through Streamlined In-Solution Digestion

**DOI:** 10.1101/2025.09.04.673655

**Authors:** Christoph Gstöttner, Tao Zhang, Sanne Pot, Lucas Hourtoulle, Katrin Heinrich, Sina Hoelterhoff, Anja Bathke, Elena Domínguez-Vega

**Affiliations:** Leiden University Medical Center, Leiden, The Netherlands; Pharma Technical Development Europe, Hoffmann-La Roche, Basel, Switzerland; Pharma Technical Development Europe, Roche Diagnostics GmbH, Penzberg, Germany

**Author notes:** The two authors contributed equally to the manuscript (Christoph Gstöttner, Tao Zhang).

## Abstract

Since the introduction of the first multidimensional liquid chromatography mass spectrometry (mD-LC-MS) approach for antibody peak characterization, multiple developments have been reported including integration of various chromatographic approaches and precise fractionation, providing high quality and reliable peptide data in a drastically decreased total analysis time. Most of these platforms rely on the use of immobilized enzyme reactors (IMERs) for digestion, limiting the applicability to few enzymes such as trypsin. Recently, the introduction of in-solution enzymatic digestion in mD-LC-MS systems has been proposed as an alternative to IMERs. Here, we make use of current innovations in 2D-LC commercial systems such as active solvent modulation valves, which permits direct mixing of the fractionated peaks with the endoprotease and reducing agent in an online manner, to integrate in-solution digestion in a straightforward manner in mD-LC-MS peak characterization platforms. Following antibody digestion, the generated peptides are automatically trapped, separated and detected by mass spectrometry. Efficient reduction and tryptic digestion was obtained using short incubation times (15 min). The approach was further expanded to alternative endoproteases such as chymotrypsin and thermolysine with other digestion specificities and allow an easy exchange between enzymes with similar buffer and digestion conditions. As a proof-of-principle that strategy was applied to achieve peptide maps from ion exchanged separated mAb peaks showing good digestion efficiency and high sequence coverages.

## Introduction

In recent years, multidimensional liquid chromatography hyphenated to mass spectrometry (mD-LC-MS) has gained attention for the automated characterization of biopharmaceuticals and many methodologies are currently in use in different biopharmaceutical labs ^1-5^. Automated mD-LC-MS approaches permit direct analysis of antibodies provided by downstream processing or formulated monoclonal antibodies (mAbs), suppressing the need of any sample preparation^3, 6-15^, but also enable online characterization of peaks separated by different chromatographic approaches. For instance, several routine chromatographic analysis such as protein A chromatography^16, 17^, ion exchange chromatography (IEX)^7, 8, 12-15^ or size exclusion chromatography (SEC)^3, 6^ have been integrated in mD-LC-

MS systems permitting in-depth characterization of separated species. The main reason for the success of these multidimensional chromatographic systems is the drastic time reduction, in particular for peak characterization from a 1D separation, due to the omission of manual fraction collection, re-analysis and overnight tryptic digestion, resulting in total analysis times in the range of hours instead of several days^7^. Furthermore, due to the fast and efficient online enzymatic digestion, the risk of introducing artificial modifications (*e*.*g*. oxidation or deamidation) is significantly reduced^8^. Another benefit of the online fractionation is that separated peaks can be more precisely fractionated than with traditional offline fraction collection.

The majority of these platforms rely on the use of immobilized enzyme reactors (IMERs) bringing some limitations in regards to enzyme applicability. Up to date, only few endoproteases such as trypsin and pepsin are commercially available in a chromatographically compatible IMER format, or can be purchased custom-made such as Lys-C^15^. This limited number of endoproteases additionally restricts the possibility to generate peptides of different length and nature, which may be required to obtain full sequence coverage of conventional mAbs, but especially in new antibody formats. Furthermore, efficient digestion often requires a previous protein reduction which introduces additional steps in the platform. In most of the cases reduction is performed in trap columns or cartridges, requiring additional pumps, followed by enzymatic digestion in IMERs^3, 7, 8, 13-15^.

Mayr et al. demonstrated recently the potential of using in-solution mAbs digestion in mD-LC-MS instead of IMERs, by presenting an automated multiple heart-cutting mD-LC design with a novel in-loop (in-solution) enzymatic heart-cut digestion between the first-dimensional column and transfer to the second dimension before MS analyses^12^. The work, furthermore demonstrated the feasibility of performing simultaneous in-solution in-loop digestion and reduction using a TCEP solution, eliminating the need for a separated reduction step. The platform comprising a pre-separation (IEC or SEC), was applied for the digestion of mAbs using trypsin as well as LysC as endoproteases or online deglycosylation using PNGaseF. However, the developed workflow was custom-made comprising several valves which are programmed to switch at the right time points limiting the implementation of the approach in different labs. Inspired by these results, we envisioned a the integration of online in-solution digestion in commercial 2D-LC systems. To this end, we make use of current innovations in 2D-LC commercial systems such as active solvent modulation (ASM) valves, which permits direct mixing of the peak cuts with the endoprotease and reducing agent in an online manner. Following digestion, the generated peptides are automatically trapped and separated on a C18 column and detected by mass spectrometry. The developed approach was further expanded to alternative endoproteases such as chymotrypsin, thermolysine and pronase with other digestion specificities and allow an easy exchange between enzymes with similar buffer and digestion conditions. The ability to easily switch between enzymes opens the possibility to close sequence coverage gaps as for instance, those encountered during a tryptic digestion, as each of these endoproteases exhibit different digestion specificity.

## Methods

### Chemicals and materials

Tris(hydroxymethyl) amino methane (TRIS) base (≥99.8%), tris(2-carboxyethyl)phosphine hydrochloride solution (TCEP, pH 7.0, 500 mM), sodium chloride (≥99.5%), sodium phosphate monobasic monohydrate (≥98%), hydrochloric acid (1N), and sodium phosphate dibasic dihydrate (99.0%), formic acid (FA) and glacial acetic acid (AA) were purchased from Sigma-Aldrich (St. Louis, MO). Trypsin, Lys-C, thermolysin, pronase, and chymotrypsin, acetonitrile (ACN, Ultra LC-MS grade) were provided by Actu-All Chemicals (Randmeer, Oss, The Netherlands). Conventional monoclonal antibody (mAb1), bispecific antibody (bsAb1) were provided by F. Hoffmann La Roche (Basel, Switzerland) and Roche Diagnostics (Penzberg, Germany). Mab1 with elevated levels of deamidation was achieved by incubation at a pH of 9.0 at 37°C for 7 days. Ultrapure water was generated from an ELGA Labwater system (Ede, the Netherlands). Prior to mD-LC-MS experiments, the formulated antibody samples were diluted to 10 mg/mL with water.

### System design and setup

The platform was built on a standard 1290 Infinity II 2D-LC system from Agilent (Agilent Technologies, Waldbronn, Germany) extended with additional modules. Overall, the system consisted of the following modules: 1260 Bio Multisampler (G5668A), one 1290 High Speed Binary Pump (G7120A), two 1260 Bio-Inert Quaternary Pump (G5654A), one DAD detector, two 1290 Multicolumn Thermostats (G7116B) configured with a 6 port/2 position valve (G5631) and a 6port/2 position external valve (G1170A). For the 2D application, the interface module consisted of a 1290 8 port/2 position valve (G4236A) and two selector valves (decks) with 6 x 120 μL loops each for multiple heart cutting (G4242A multiple heart cutting kit). For the protein digestion an additional sample loop (Agilent 5500-1456, Prep LC capillary ST, 0.6×900 mm) was implemented in the system between the ASM valve and the C18 column. The system was controlled by two OpenLab software packages, one for the two quaternary pumps, the two ovens, and the external valve and the second for the remaining modules. The “remote” start signals are delivered by a contact closure board integrated in the autosampler. The mD-LC was directly connected to a QTOF mass spectrometer (Bruker Daltonics, Bremen, Germany), equipped with an electrospray ionization (ESI) ion source.

#### 1. First dimension: Ion exchange separation and fractionation

In the first dimension, charged antibody variants were separated by IEC separation on a 4 x 250 mm ProPac WCX-10 analytical cation exchange column (10 µm, Dionex, Thermo Scientific, Massachusetts, USA). A gradient from 15% to 50% B in 30 min using 10 mM sodium phosphate (pH 7.5) as solvent A and 10 mM sodium phosphate buffer, 0.1 M NaCl (pH 7.5) as solvent B was applied for mAb1 (**Table S1**). For bsAb1 and deamidated mAb1, a gradient from 5% to 55% B in 25 min with 10 mM sodium phosphate (pH 7.0) as solvent A and 10 mM sodium phosphate buffer, 0.1 M NaCl (pH 7.0) as solvent B was employed (**Table S2**). The flow rate was 800 µL/min. The IEC separation was performed at room temperature with UV detection at 280 nm. For each analysis, 100 μg of mAb were injected. Fractionation of mAbs was performed based on heart-cutting and fractions of interest which were stored in one of the twelve 120 µL loops of decks A and B.

#### 2. Protein reduction and enzymatic digestion using active solvent modulation

Immediately after the first fraction was collected in one of the loops of deck A or B, the collected fraction was flushed out of the loop by the second dimension quaternary pump at a flow rate of 60 µL/min for 4 min. Due to the usage of an active solvent modulation (ASM) valve and chosen dilution factor of 1:1, the fraction stored in one of the twelve fraction collection loops was directly diluted with the digestion buffer containing the reducing agent, the trypsin solution and the digestion buffer. Mobile phase A was composed of 50 mM Tris-HCl buffer at pH 7.5 with 2 mM TCEP and mobile phase B containing 10mM Tris-HCl, pH 7.5 and enzyme at concentration of 4µg/mL for Lys-C, pronase and thermolysin. The used mobile phases containing the digestion enzymes were prepared freshly every day, while the mobile phase A was refreshed every weekly. In the case of trypsin and chymotrypsin, mobile phase A consisted of 50 mM Tris-HCl buffer at pH 9.3 and 8.5 with 2 mM TCEP, respectively. Mobile phase B contained the endoproteases at a concentration of 4 µg/mL trypsin or chymotrypsin in 20 mM acetic acid to prevent self-digestion. Detailed enzyme buffer conditions are summarized in **Table S3**. After mixing the sample with the reducing agent and the endoprotease solution, the mixture was incubated in a large sample loop (Agilent 5500-1456, Prep LC capillary ST, 320 µL, 0.6×900 mm) for 15 min at a low flow rate of 1 µL/min to achieve subsequent antibody reduction and enzymatic digestion of each individual fraction. The sample loop was installed in a column oven at a temperature of 37 □ with the exception of thermolysin and chymotrypsin, which were incubated at 85 □ and 50 □, respectively. After reduction and digestion, the generated peptides were flushed to the RP column with 50% mobile phase B at a flow rate of 100 µL/min for 6 min where the peptides are trapped. To prevent carryover the large sample loop was flushed with 20% ACN (Mobile phase C) for 4 min at a flow rate of 500 µL/min followed by a flush with the reduction/digestion solution for the next antibody fraction analysis (50% mobile phase B) for 10 min at a flow rate of 60 µL/min. During the cleaning step the valve located after the sample loop was switched to waste position. Detailed gradients and valve positions are summarized in **Table S4**.

#### 3. Second dimension: peptide mapping

The generated peptides were separated on a RP XSelect CSH C18 column (130Å, 3.5 µm, 2.1 mm X 100 mm column Waters, Mildford, MA). The mobile phases were composed of water (A) and ACN (B) containing both 0.1% FA. The flow rate was set to 400 µL/min and the column temperature was 60°C. The trapped peptides were first washed to prevent contamination of the MS system by the Tris containing digestion buffer, using 1% mobile phase B for 1 min. Following the cleaning step, the RP-C18 column was set in-line with the MS via an external switch valve. The peptides were subsequently separated using a multiple step gradient from 1% to 15% B in 6 min, from 15% to 19% B in 9 min, from 19% to 35% in 7 min and from 35% to 70% solvent B in 10 min via the binary 2D pump and analyzed by ESI-qTOF. At the end of the gradient the column was flushed with 100% Mobile Phase B for 3 min. After the elution step was completed the large sample loop was switched again in-line with the RP column to analyze the next fraction. The gradient details and valve switching times are shown in **Table S5**. The entire process is depicted schematically in **Figure 1**.

**Figure 1.**
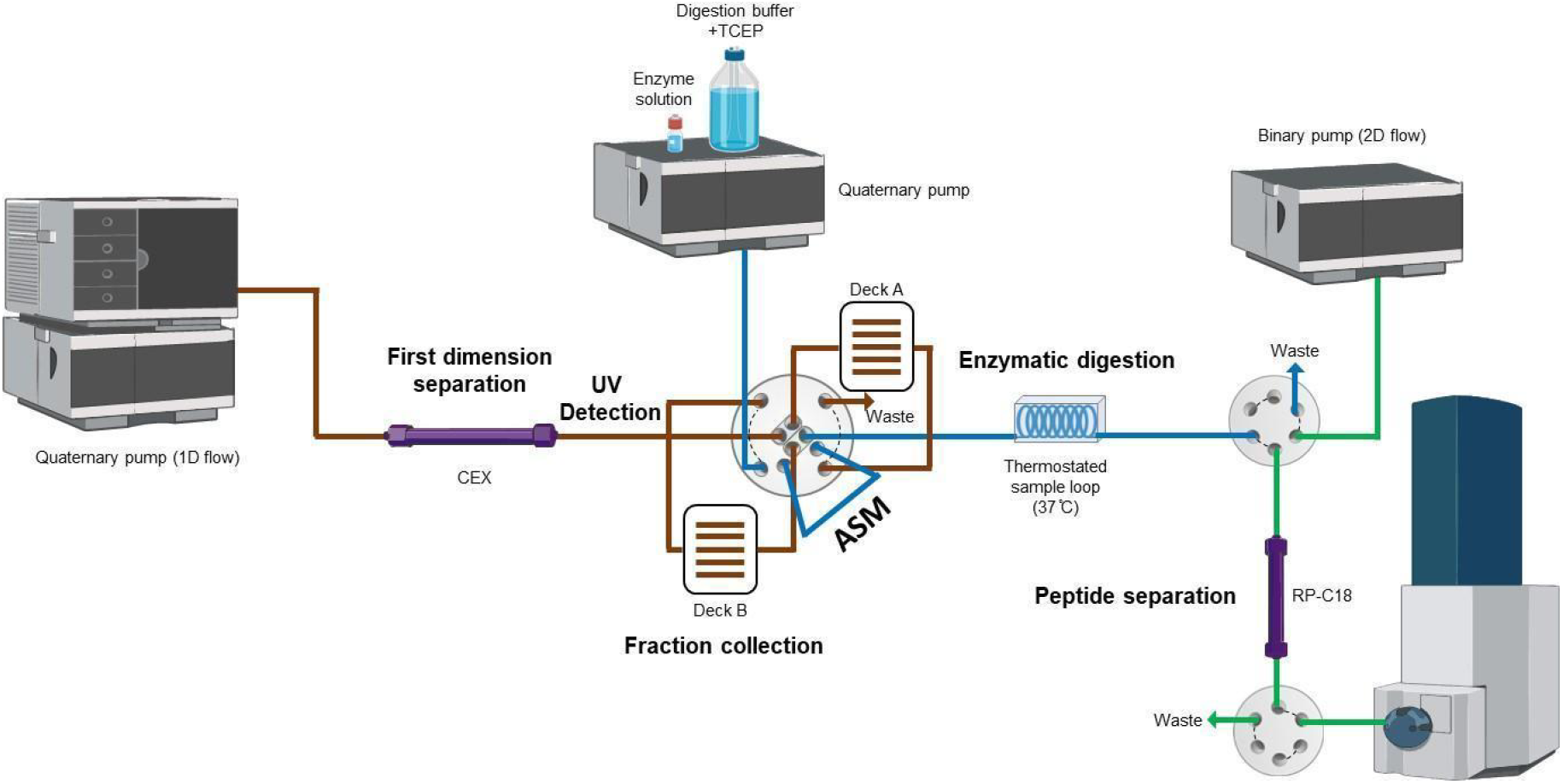
Schematic representation of the in-solution online digestion multidimensional LC-MS platform.

### LC-MS data collection and analysis

The MS analysis was performed on an Impact II ESI qTOF (Bruker, Bremen, Germany). Data was acquired using the HyStar acquisition software from Bruker Daltonics. For peptide acquisition the following parameters were employed: *m/z* range from 150 to 2500, transfer time of 130 µs, pre pulse storage time of 15 µs, collision energy was 7 eV, 500 V end plate offset, 4500 V capillary voltage, 5.5 L/min dry gas flow rate, 180□ dry gas temperature and 2 bar nebulizer gas pressure. MS data processing was manually performed using DataAnalysis (Bruker Daltonics) and Skyline 20.2 software (Mac-Coss Lab, U Washington). The Skyline software settings were as following: Number of allowed missed cleavages up to 1 (in case of sequence coverage calculation) and 3 (in case of missed cleavage detection), precursor charges were set to 2, 3 and 4 and for the MS1 filtering TOF was chosen as a mass analyzer with a resolving power of 30.000. The sequence coverage was calculated manually by checking the corresponding peptides in Skyline.

## Results and discussion

### Online in-solution digestion platform development

In this work, we aimed to develop an in-solution online digestion strategy for mD-LC-MS peptide map analysis of mAbs using commercial 2D-LC instrumentation. The platform should enable pre-separation (e.g. by IEC), peak fractionation, in-solution reduction and enzymatic digestion as well as a C18 RP separation of the peptides followed by MS detection (**Figure 1**). In this work, we selected IEC as first dimension separation due to its extensive usage for routine mAb characterization. After separation, five 120 µL fractions were automatically fractionated using the multiple heart-cutting mode and temporarily stored in the loop. To enable an efficient online in-solution digestion few aspects need to be considered. First, the mAb species fractionated in the loops are dissolved in the IEC mobile phase while the digestion often requires different buffer environments. Therefore, we make use of the active solvent modulation (ASM) valve which enables efficient mixing of fractions with the digestion solutions. Importantly, endoproteases such as trypsin and chymotrypsin may undergo autoproteolysis when stored in the digestion buffer. Therefore, two independent solutions were employed containing (A) the enzyme in acetic acid solutions and (B) the digestion buffer containing the reduction reagent (TCEP) which were mixed with the quaternary pump used to deliver the solution to the ASM valve. Using the ASM functionality with a dilution factor of 1:1 the collected fractions were mixed with TCEP and the endoprotease resulting in a final concentration of 0.5 mM TCEP and 1ug/mL endoprotease after mixing. To allow in-solution digestion a large sample loop (∼ 250 µL) was added after the ASM allowing and inserted in a thermostated column compartment to allow incubation for online reduction and enzymatic digestion at the desired incubation temperature. After reduction and digestion the generated peptides were analyzed by C18 LC-MS. During C18 peptide mapping, the sample loop used for digestion was extensively washed using 2 mL 50 mM Tris-HCl buffer pH 7.5 to significantly decrease any carryover, following a flush with the digestion buffer containing TCEP to equilibrate for the next analysis. Using the proposed configuration only three pumps were required (one for the 1D gradient, one for the enzyme and digestion buffer and one for the peptide separation), reducing thereby costs but also the total analysis time compared to previously mentioned mD-LC-MS platforms using IMERs, which require an additional pump for the reduction step.

### Online in-solution mAb digestion with trypsin

In-solution endoprotease digestion occurs in the sample loop following mixing through the ASM valve. To determine the optimal digestion conditions we assessed different digestion/incubation conditions including enzyme concentration, reaction time and digestion buffer pH. Trypsin was employed for the optimization as it is one of the most commonly employed endoproteases. To prevent self-cleavage of trypsin the enzyme was dissolved in 20 mM acetic acid at a concentration of 4 µg/mL (mobile phase B). Trypsin was mixed with the digestion buffer in the quaternary pump prior to incubation with the collected fractions. The digestion buffer which provided adequate pH consisted of 50 mM Tris-HCl buffer at pH 9.3. After mixing the digestion buffer at pH 9.3 with the enzyme solution in 20 mM acetic acid, the resulting pH was 8.2, which is within the optimal pH range for trypsin digestion. In addition, as suggested by Mayr et al., 2 mM TCEP (final concentration 0.5mM TCEP) was added to the digestion buffer to simultaneously enable reduction of the disulfide bonds (**Figure S1**). This enabled the elimination of the additional reduction step in the trap column required in previous mD-LC-MS platforms incorporating IMERs, leading to a simpler setup. After digestion, the peptides were flushed from the sample loop to the RP column using the digestion buffer followed by a rinse with 1% ACN containing 0.1% FA to remove the salts.

The effect of incubation time (30 min, 15 min and 5 min) on digestion efficiency and sequence coverage was evaluated. As shown in **Figure 2**, both 30 min and 15 min reaction times showed efficient trypsin digestion and high sequence coverage (100% for LC and 99.1% for HC). Digging into the generated peptides, the same number and nature of peptides were detected in MS using both, 15 or 30 min, incubation times (**Figure S2, Table S6**). Furthermore, only minor amounts of light chain (LC) were observed indicating an efficient automated enzymatic digestion (**Figure 2B**). The levels of LC were slightly lower in the case of 30 min digestion time they but did not influence the sequence coverage. Shorter reaction times (5 min) were evaluated, however, resulted in a significant decrease in signal intensity and sequence coverage of the heavy chain from 99.1% to 94.2%. Regarding miscleavages, overall similar number of miscleavages were observed independently of the digestion time (**Table S6**). Additionally, five exemplary polar peptides were evaluated in terms of retention time and areas, providing very comparable results for 30 min and 15 min incubation time (**Figure S3**). For 5 min we observed a significant decrease in the area of the selected polar peptides including a loss of the peptide VTITCR.

**Figure 2.**
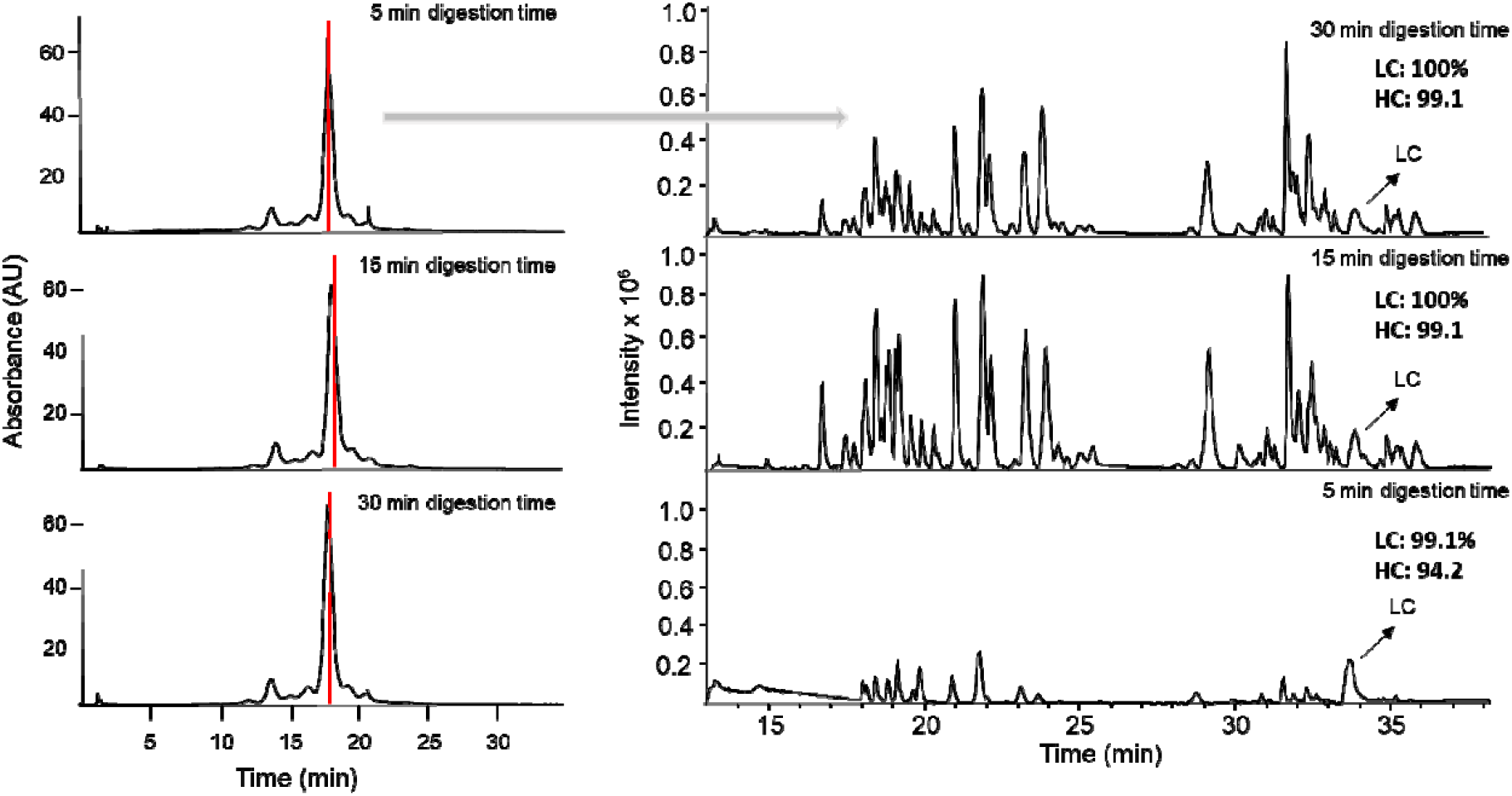
Influence of incubation time on digestion efficiency of mAb1 using trypsin online in-solution for 5 min, 15 min or 30 min. A) CEX separation of mAb1 in first dimension highlighting the cut in the main peak used for further digestion. B) Base peak chromatograms (BPC) of the peptide map obtained after online in-solution digestion for the selected cut. The BPCs were aligned to allow an easier comparison between the different incubation times. The sequence coverage is shown in bold for the LC and HC. *LC, light chain.

Additionally two different concentrations of trypsin concentration were evaluated in the online in-solution digestion (1 and 2 µg/mL trypsin after mixing) (**Figure S4**). Lower signal intensity was observed when mAb1 was digested with 1 µg/mL and therefore a concentration of 2 µg/mL trypsin was selected for the experiment. Higher amounts were not evaluated as good sensitivity of the peptides was observed while maintaining a good cost-efficiency of the platform (2 µg of trypsin per run).

Next, the approach was evaluated in a multiple cut environment (**Figure S5**). In the IEC separation the deamidated mAb is eluting at 14 min^14^. Therefore, the peak corresponding to the deamidated antibody was online fractionated, digested and analyzed (fraction 1 in **Figure S5**) followed by the same process for the main peak (fraction 2 in **Figure S5**). The peptide map obtained for fraction 1 indeed showed high levels of deamidation, while the peptides generated from fraction 2 (main peak) corresponded to unmodified peptides.

The reproducibility of the digestion was evaluated within one analysis by cutting three fractions within the main peak and between analysis on three different days. **Figure S6** shows no difference in retention time as well as charged variant separation between different IEC injections, which is a prerequisite to allow a comparison of digestion efficiency between different runs. Reproducible peptide maps and sequence coverage (LC, 100% and HC, 98.1%) were observed across all three injections within the same analysis but also between three analysis on different days (**Figure S7**). Additionally the retention times of five polar peptides were compared resulting in RSD values between 0 and 0.04% within the same sequence and between 0.16 and 0.2% between different sequences and days demonstrating the good reproducibility of the system (**Table S7**).

### Increasing applicability of in-solution online digestion to a broad range of endoproteases

mD-LC-MS approaches for peptide mapping typically rely on IMERs, however, commercially available IMERs are limited to trypsin or pepsin, restricting the applicability of this approach to very few proteases, which can result in gaps in the sequence coverage. Alternative proteases are often required in biopharmaceutical characterization to ensure good sequence coverage of different antibodies and novel formats, to fill tryptic gaps or to improve separation of certain PTMs. The proposed online in-solution digestion strategy eliminates the need for an immobilized column, allowing to expand the digestion to other endoproteases.

To further demonstrate the capabilities of this platform for alternative enzymatic digestions, we evaluated several widely used endoproteases for bottom-up proteomics, including Lys-C, thermolysin, chymotrypsin and pronase. Given the different optimal pH and temperature ranges of these enzymes, 50 mM Tris buffer was adjusted to the required pH taking also into account the final pH after mixing with the enzyme solution (**Table S3**). The temperature was easily adjusted by modifying the temperature of the column compartment in which the digestion loop was integrated. Most of the enzymes were employed with a final pH of 7.5. In contrast, chymotrypsin digestion was performed under slightly modified conditions, using a higher pH of 7.8 and an elevated temperature of 50°C, as suggested by the provider. For thermolysin the incubation temperature was set at 85°C following the recommendations for the digestion. **Figure 3** shows the BPC obtained for LysC, Chymotrypsin and thermolysin after online in-solution digestion for 15 min. LysC showed a very high digestion efficiency with a sequence coverage of 100% for the light chain and 99.1% for the heavy chain. In case of chymotrypsin the light chain sequence coverage was 100% whereas for the heavy chain the sequence coverage was 99.2%. Thermolysine showed a slightly decreased sequence coverage of 93.0% for the light chain and 86.2% due to smaller peptides or even single amino acids generated, which are not retained on the C18 column. Yet, thermolysin can be useful endoprotease to cover trypsin gaps such as the VEPK peptide not detected after tryptic digestion (**Figure 3B**).

**Figure 3.**
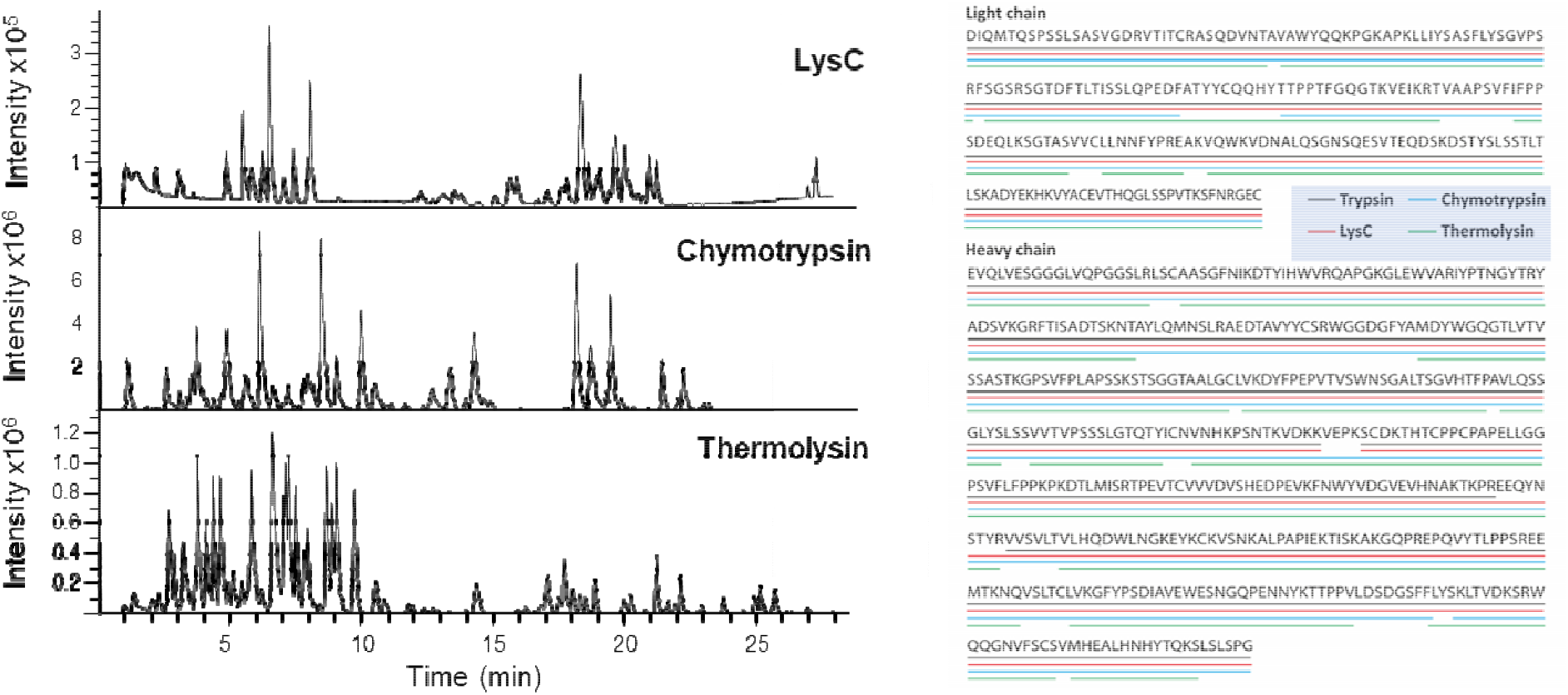
A) BPC and B) sequence coverage obtained for mAb1 after online in-solution digestion using different endoproteases. The sequence coverage was manually checked in the Skyline software and is depicted by colored lines: grey – trypsin, blue – chymotrypsin, red – lysC and green – thermolysin.

Pronase is an enzyme of broad specificity used for protein digestion to generate smaller peptides suitable for MS analysis of complex proteins. To achieve efficient digestion with pronase a longer incubation time (i.e. 30 min) was necessary. As shown in **Figure S8**, good peptide profile was also observed for pronase, however, still some remaining levels of undigested LC and HC were observed even after 30 min incubation. Yet, the enzyme could be useful for the characterization of complex Ab formats or other proteomic applications either alone, or in combination with other enzymes.

### Online in-solution digestion in mD-LC-MS of bispecific antibodies

To demonstrate the capabilities of the mD-LC-MS set-up, the method was applied to the analysis of more complex antibodies such as bispecific antibodies. For bsAb1 a different gradient and pH of mobile phase A and B were employed (**Table S2**) due to the low retention observed in IEC with the conditions used for mAb1. As shown in **Figure 4**, different separated peaks were observed after IEC separation of bsAb1. The digestion of the main peak using trypsin and Lys-C, respectively, was evaluated using the setting described in the materials and methods (**Figure S9**). In both cases good digestion of bsAb1 was obtained indicating that differences in formulation do not interfere with the analysis. Furthermore, five fractions of bsAb1, corresponding to three different IEC separated peaks at retention times 16.53, 18.50, 18,73, 20.25, and 20.62 min were collected following online digestion with Lys-C. All fractions were efficiently digested independently of their elution time and concentration with sequence coverages between 97.2-100% (**Figure 4B**) and number of miscleavages (**Table S8**). Furthermore, the retention time and areas of four representative peptides were evaluated (**Figure S10**) showing again good repeatability demonstrating the applicability of this approach for new antibody-derived therapeutics.

**Figure 4.**
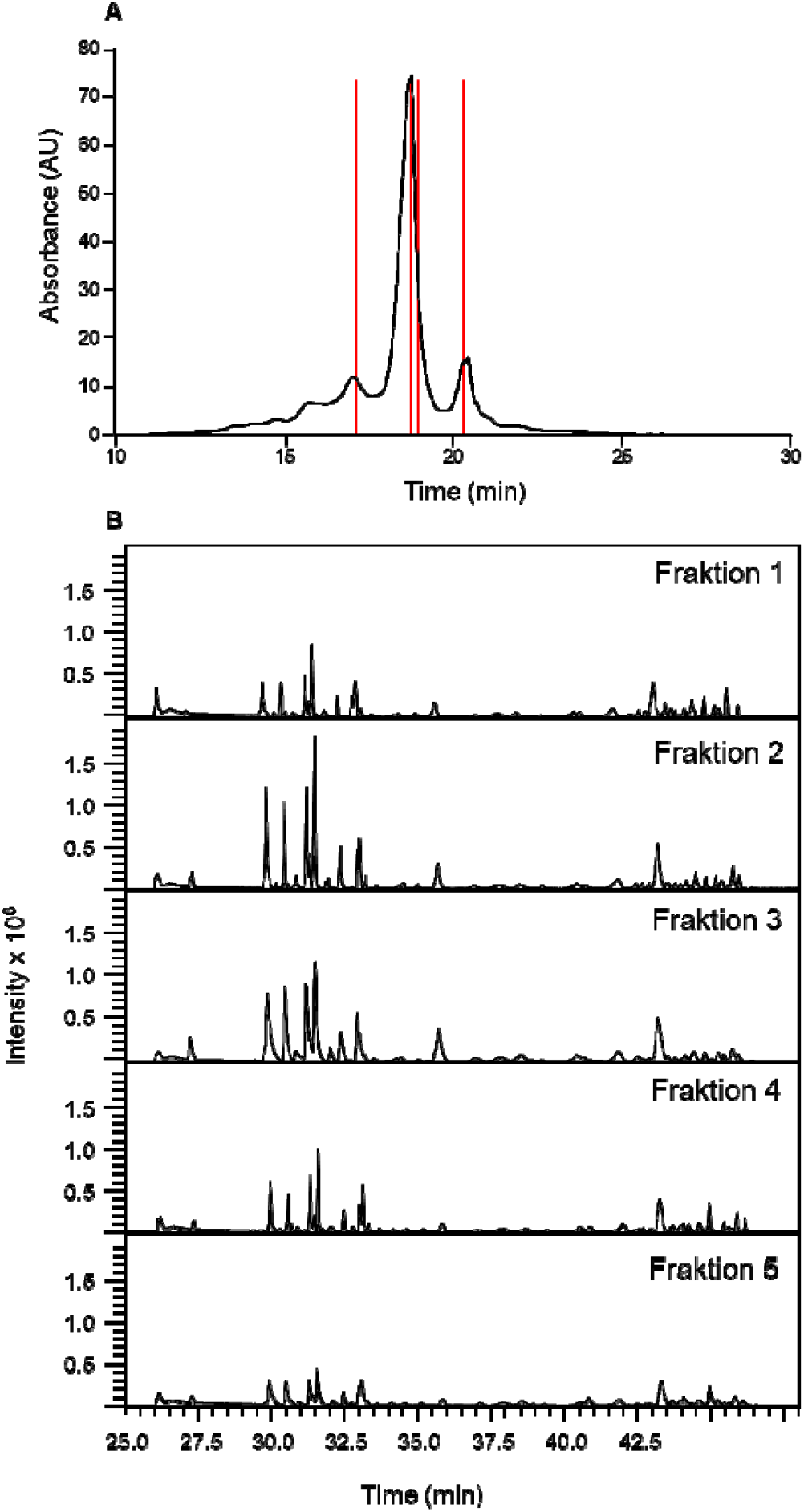
A) IEC chromatogram of bsAb1 and fractions collected for further online in-solution digestion. Fractions are depicted in gray. B) BPC of the peptide map obtained after online in-solution digestion of different fractions of bsAb1.

## Conclusions

In this work, we developed an in-solution online digestion strategy making use of the ASM valve present in commercial 2D-systems for peptide map analysis of mAbs, enabling pre-separation, peak fractionation, in-solution reduction and enzymatic digestion as well as a C18 RP-LC-MS analysis of the peptides. The integration of peak fractionation with online in-solution reduction and digestion resulted in a simplified platform compared to our previous mD-LC-MS setups requiring fewer pumps and reducing the total analysis time^7, 12^. Efficient and fully automated enzymatic digestion of antibodies was achieved, supported by the high sequence coverage as well as minimal detection of undigested material. The platform was further evaluated using a wide range of alternative proteases demonstrating effective digestion with minimal adjustments in digestion buffer composition or incubation temperature. This expanded enzymatic flexibility enhances the versatility of peptide mapping workflows, enabling tailored digestion strategies beyond conventional trypsin-based methods. By obtaining orthogonal proteolytic cleavages, tryptic gaps or complex proteins can now be addressed using complementary enzymes. The online in-solution digestion also opens the possibility for combining endoproteases with different digestion specificity in case one endoprotease does not generate adequate peptides and will be evaluated in future studies. The developed platform represents thus an important addition to the toolbox of automated biopharmaceutical digestion platforms.

## Author contributions

*CG and TZ contributed equally the manuscript. CG, TZ and SP performed the experiments. CG, TZ, LH, KH, SH, AB, and ED conceptually designed the work. CG, TZ and ED wrote the manuscript. All authors have given approval to the final version of the manuscript.

## Notes

The authors declare no competing financial interest.

## Table of Content

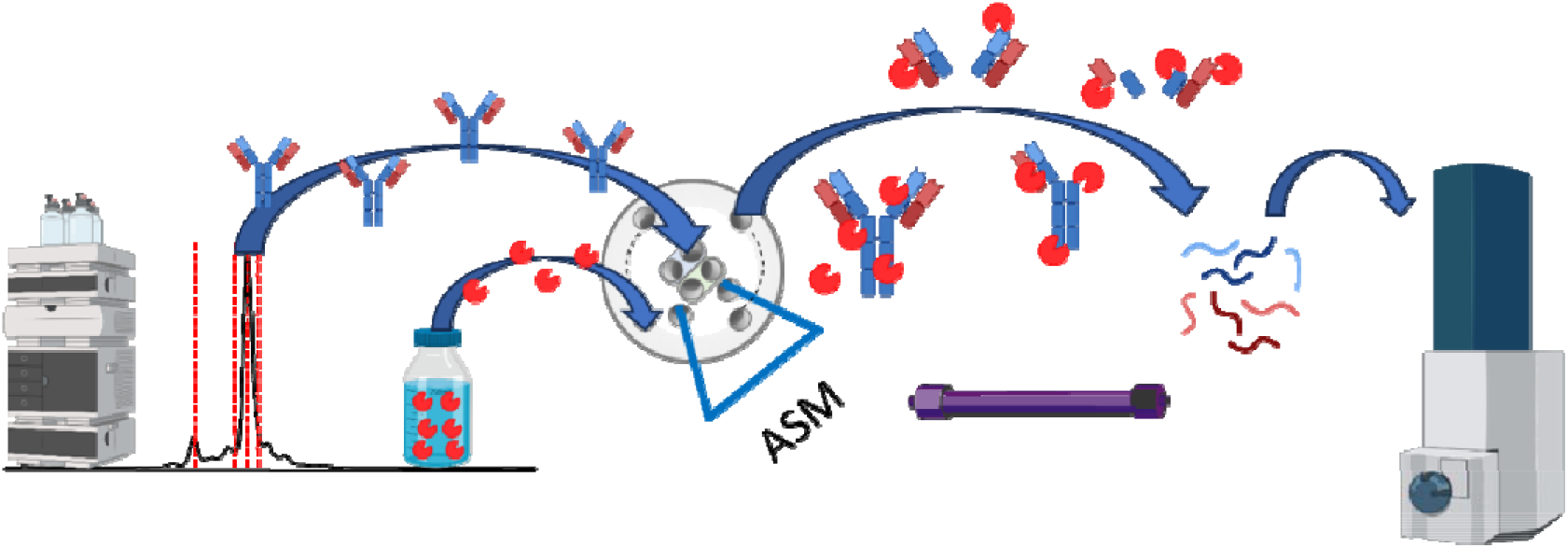

