## Supporting information for "Increasing Applicability of Automated mD-LC-MS Peptide Mapping for Biopharmaceuticals through Streamlined In-Solution Digestion"

**Supplementary Figures**

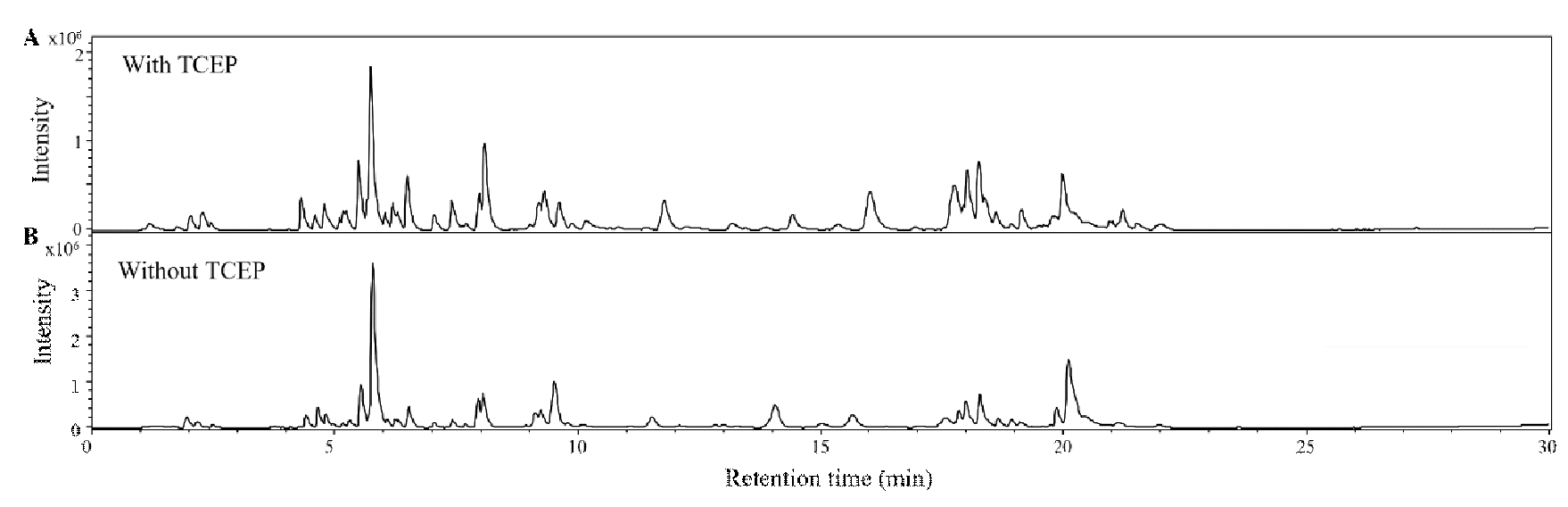

**Figure S1.** Base peak chromatograms (BPC) of the peptide map obtained after online in-solution digestion for the main peak of mAb1 after IEC with and without TCEP reduction.

.

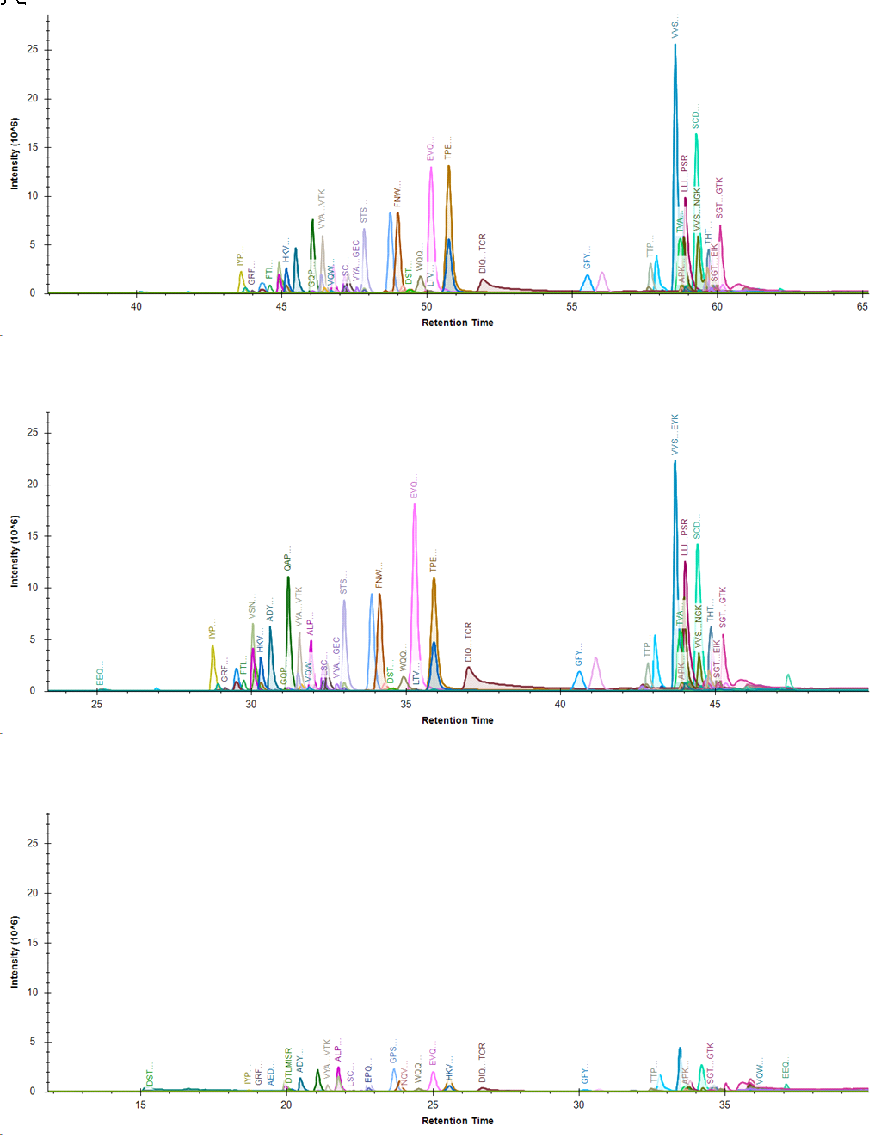

**Figure S2**. Extracted Ion Chromatograms (EIC) of the peptide map obtained after online in-solution digestion for the main peak of mAb1 after IEC using different incubation times. The different time axis is a consequence of the different incubation times used.

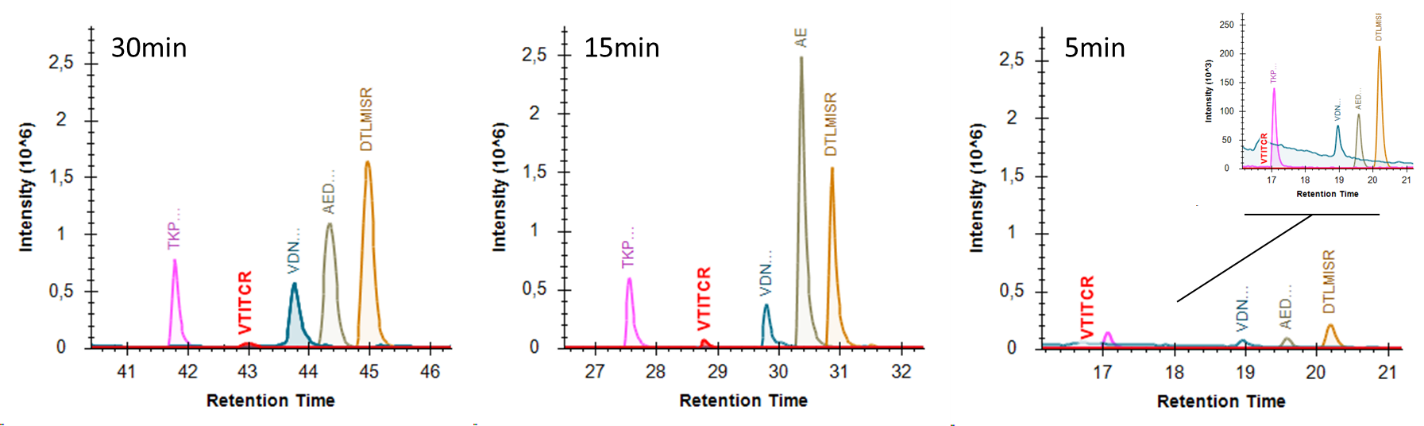

**Figure S3**. Extracted Ion Chromatograms (EIC) of the five polar peptides obtained after online in-solution digestion for the main peak of mAb1 after IEC using different incubation times (5min, 15 min and 30min). The different time axis is a consequence of the different incubation times used.

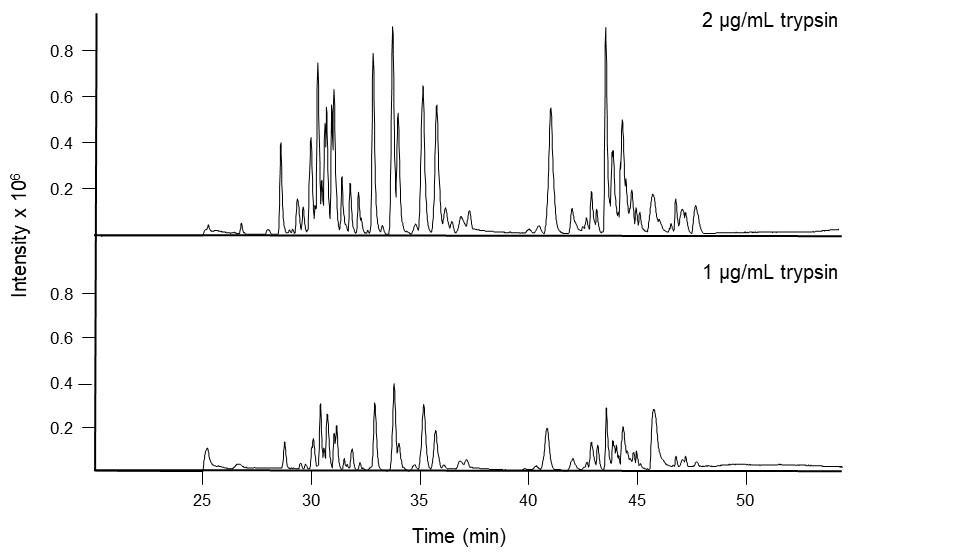

**Figure S4.** 2D base peak chromatograms (BPC) of the peptide map obtained after online in-solution digestion for the main peak of mAb1 after IEC using 2 µg/mL or 1 µg/mL trypsin

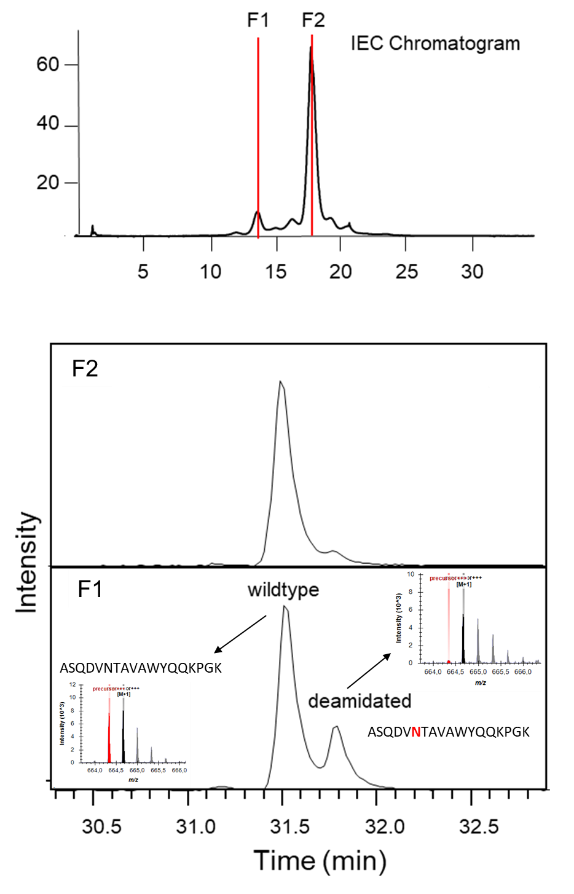

**Figure S5**. A) CEX separation of stressed mAb1 highlighting the cut in the main peak used for further digestion. B) Extracted ion chromatogram of one of the peptides exhibiting a deamidation spot after on-line in-solution digestion for the selected cut (F1 or F2).

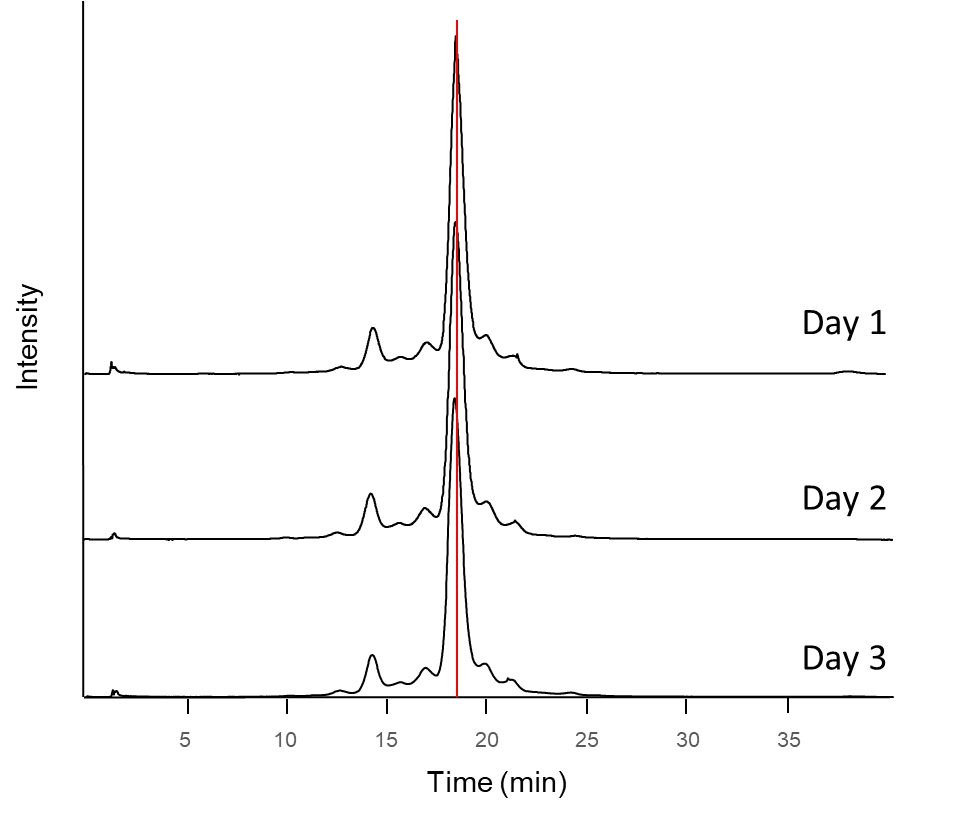

**Figure S6.** IEC chromatogram of mAb1 in three independent days. Fractions are depicted in red.

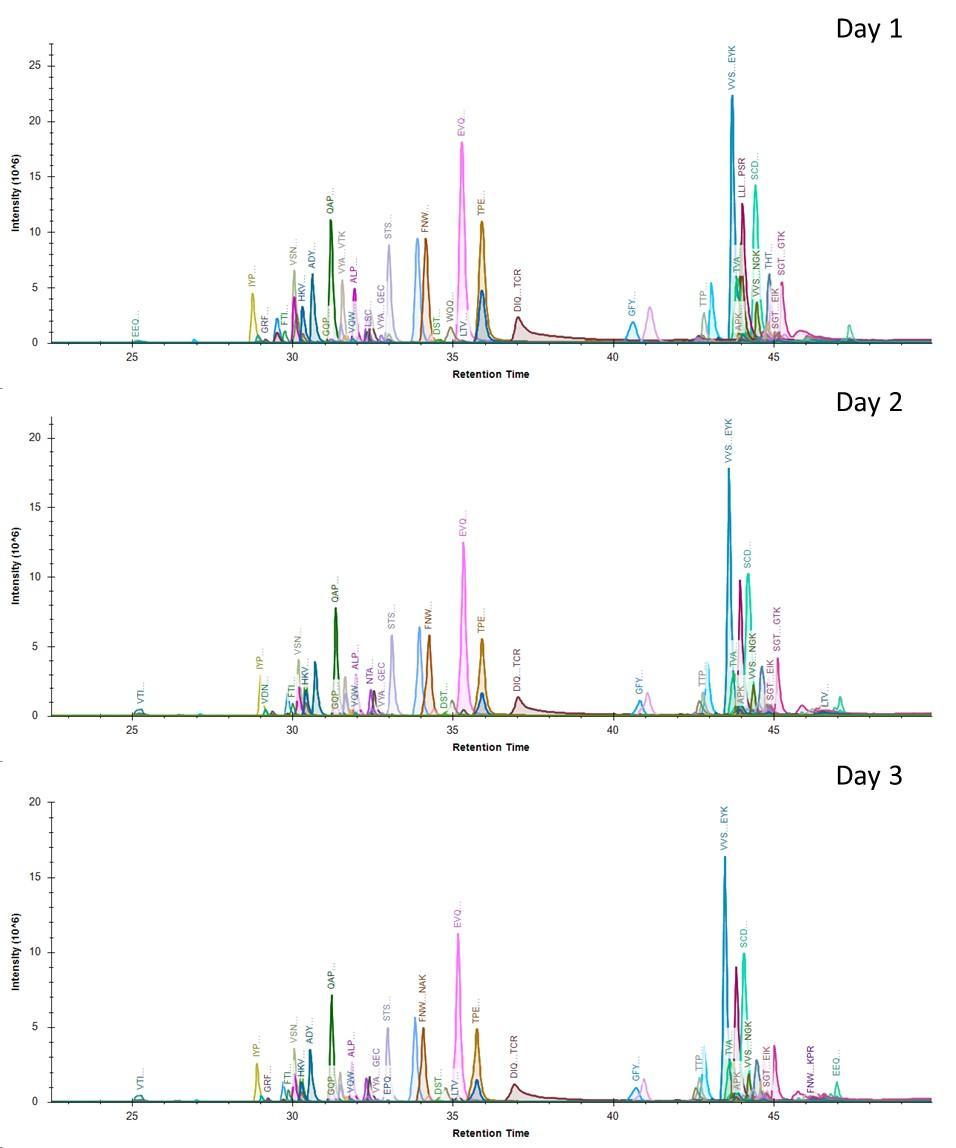

**Figure S7**. 2D base peak chromatograms (BPC) of the peptide map obtained after online in-solution digestion for the main peak of mAb1 after IEC in three independent days.

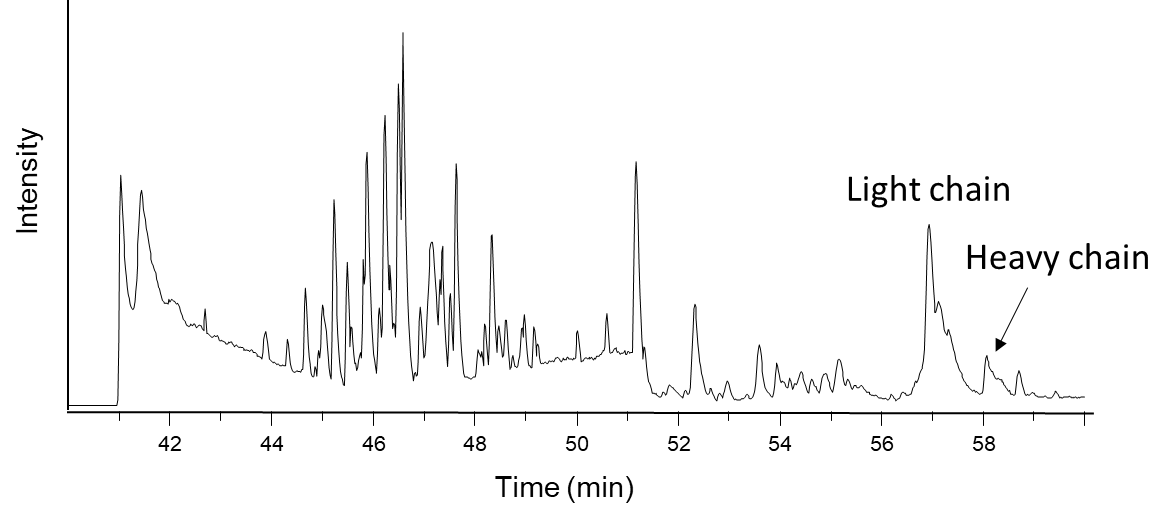

**Figure S8**. BPC of the peptide map obtained after online in-solution digestion of the main peak of mAb1 after IEC using Pronase.

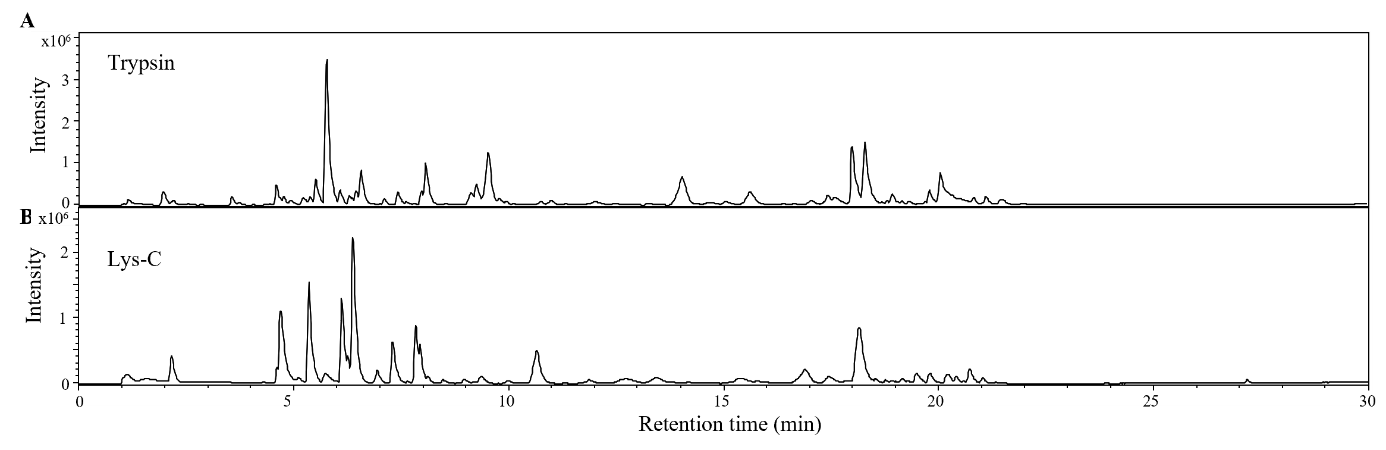

**Figure S9**. BPC of the peptide map obtained after online in-solution digestion of the main fraction of bsAb1 after IEC (fraction 4) using trypsin (A) or Lys-C (B).

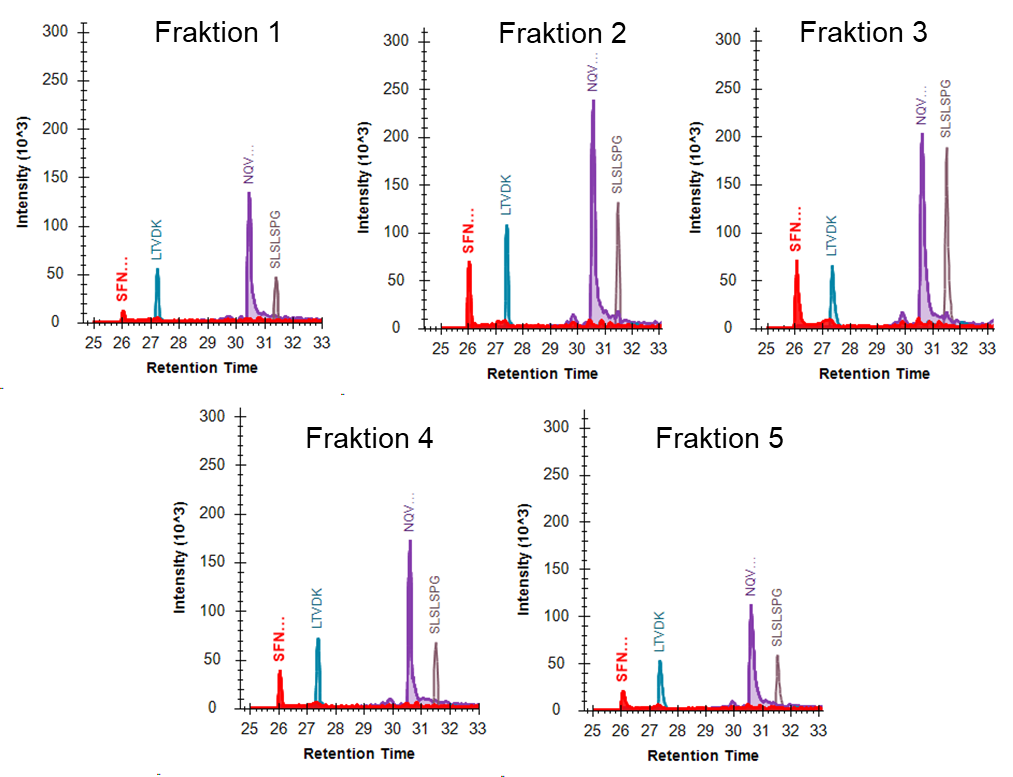

**Figure S10**. Extracted Ion Chromatograms (EIC) of the four polar peptides obtained after online LysC in-solution digestion for five different IEC fractions of bsAb1.
